# A blueprint for the implementation of a validated approach for the detection of SARS-Cov2 in clinical samples in academic facilities

**DOI:** 10.1101/2020.04.14.041319

**Authors:** Sushmita Sridhar, Sally Forrest, Iain Kean, Jamie Young, Josefin Bartholdson Scott, Mailis Maes, Joana Pereira-Dias, Surendra Parmar, Matthew Routledge, Lucy Rivett, Gordon Dougan, Michael Weekes, Martin Curran, Ian Goodfellow, Stephen Baker

## Abstract

The COVID-19 pandemic is expanding at an unprecedented rate. As a result, diagnostic services are stretched to their limit, and there is a clear need for the provision of additional diagnostic capacity. Academic laboratories, many of which are closed due to governmental lockdowns, may be in a position to support local screening capacity by adapting their current laboratory practices. Here, we describe the process of developing a SARS-Cov2 diagnostic workflow in a conventional academic Containment Level 2 (CL2) laboratory. Our outline includes simple SARS-Cov2 deactivation upon contact, the methods for a quantitative real-time reverse transcriptase PCR (qRT-PCR) detecting SARS-Cov2, a description of process establishment and validation, and some considerations for establishing a similar workflow elsewhere. This was achieved under challenging circumstances through the collaborative efforts of scientists, clinical staff, and diagnostic staff to mitigate to the ongoing crisis. Within 14 days, we created a validated COVID-19 diagnostics service for healthcare workers in our local hospital. The described methods are not exhaustive, but we hope may offer support to other academic groups aiming to set up something comparable in a short time frame.

## Introduction

SARS-Cov2, the viral agent of COVID-19, is a recent introduction into the human population, and the disease epidemic is expanding nationally (within the UK) and internationally at an unprecedented rate [1,2]. As a result, national diagnostic services are stretched to and there is a need for alternative laboratory facilities to provide additional diagnostic capacity. The screening of asymptomatic individuals and healthcare workers (HCWs) will be key for controlling the epidemic and also to ensure HCW are a) working in safe conditions with functional personal protective equipment (PPE), b) not transmitting SARS-Cov2 to vulnerable patients on wards, and c) can return to work if they are not actively infected [3]. However, in almost all cases, embedding a rapid testing workflow for screening asymptomatic individuals and HCWs within the current hospital structure would add additional pressure upon an already overstretched diagnostic service.

The UK government and other organisations have recognised the critical role of additional screening [4]. For example, the UK has recently initiated the establishment of national testing centres for HCWs; however, there is a need for guidance on how such systems be standardised or scaled. The expected turnaround times from sampling collection to results being reported back to the affected HCW is also a key issue. Whilst central screening facilities will ultimately be beneficial in curtailing the epidemic, smaller academic and non-academic laboratories can (and should) contribute to these efforts. Critically, local facilities (academic laboratories in proximity to or within healthcare facilities) can frequently provide a quicker turnaround time than larger remote facilities due to simpler sampling and shipping logistics or even overstretched, onsite diagnostic laboratories; current turnaround time is typically >48 hours from sample being taken to provision of result. Such delays can have a negative impact on healthcare provision, for example staffing levels may be strained due to HCWs isolating as a result of a respiratory illnesses other than COVID-19. Therefore, there is an urgent unmet need to establish local screening workflows for HCWs and those working in essential service industries.

One of the key limitations associated with the expansion of diagnostics to tackle the COVID-19 outbreak is the availability of protocols, or a scheme, that can be used in suitable laboratory for detecting SARS-Cov2 in suitable clinical samples. In the given circumstances, a robust diagnostic test for COVID-19 should be able to generate a result rapidly, but also maintain a high level of reproducibility, specificity, and sensitivity [5]. The test also needs to be conducted on an easily accessible clinical sample, such as a nose/mouth swab, urine, or blood. Such tests can be based upon a direct amplification assay for a component of the viral genome, a suitable biomarker or metabolic signature, or the measurement of an indicative acute antibody response. Given a paucity of reliable alternatives, a PCR based approach is currently the most suitable and scalable model, whilst providing an acceptable compromise between turnaround time and accuracy.

The key issues for rapidly establishing a new diagnostic testing platform are sampling, safety, reagents, cleanliness, methodology, and reporting. Early indications for COVID-19 infections is that there are relatively high titres of virus in the respiratory tract, possibly in the gastrointestinal tract, but lower concentrations in blood [6]. Consequently, oral/nasal swabs are widely accepted as the optimal clinical sample for HCWs and others who are likely to have a higher occupational exposure risk to the virus through their work. These swabs need to be handled safely, so the use of a sampling method that inactivates the virus rapidly is essential to protect the sampler and those handling the sample, including couriers, and laboratory staff [7]. The availability and the expense of reagents required for an effective testing programme at a specific scale (e.g. hospital, company, or local community) is critical giving the ongoing demand for specific kits. Tests that require expensive mainstream reagents, or those in short supply, should be avoided where possible. PCR diagnostics are prone to issues with contamination and appropriate workflow and strict sample/reagent segregation needs to be adopted, which may be problematic in some setting. Methodologies and equipment are variable, but every attempt should be made to ensure the tests are performed using a standardised and validated test with appropriate controls. Lastly, the resulting data needs to be authenticated by a qualified individual and reported in a timely fashion through an existing and official reporting system, whilst at the same time ensuring patient confidentiality.

Here we describe our experience in establishing a COVID-19 diagnostics laboratory in an academic containment level 2 (BSL2) research facility (UK) in which we validated and established a real-time PCR workflow to detect SARS-CoV2 in nasal/oral from HCWs. We developed an assay and workflow over eight working days (set-up to validation to screening) that can produce a quantitative diagnostic result ~4 hours after swabbing.

## Methods

### Swabbing

For the swabbing of known COVID-19 patients and HCWs we developed a kit that can be easily assembled and provided in bulk. The kit contained; swabbing instructions, an individually packed sterile swab that can be broken (VWR), a labelled sample tube, and gloves (see protocol 1).

The instructions indicate the individual to put on the gloves, remove the sterile swab from the packet and swab the rear of the mouth, the back of throat, and then each nasal cavity (one swab, four sites). The swab is placed into the labelled sample tube (4ml long necked externally threaded cryovials (Nunc 379146) to avoid aerosols) and the end is submerged in 500μl transport medium/lysis buffer (4M guanidine thiocyanate (Merck) in 25 mM Tris-HCl, 0.5% β-mercaptoethanol (Sigma), and carrier RNA (100μl of 1μg/μl stock; Qiagen). The swab is snapped carefully to avoid disturbing the buffer and the cap is place back onto tube containing the buffer and swab and tightened. The tube is gently agitated to ensure even distribution of lysis buffer and labelled with an ethanol resistant pen. The outside of the tube is sprayed with 80% ethanol, placed into a zip lock bag (Onecall) and sealed. One glove is removed, and the zip lock bag is sprayed with 80% ethanol while being held in gloved hand and then passed to a clean hand. The sealed bag is place in a secure biohazard labelled container for dispatch to a certified BSL2 laboratory.

### Nucleic acid extraction

The combination of 4M guanidine thiocyanate and 0.5% β-mercaptoethanol should ensure complete lysis and deactivation of the virus but to ensure additional safety the samples are received and unpacked in a class II microbiological safety cabinet (MSC) (see protocol 2 and buffer preparation) [8]. Notably, whilst this process should be conducted in a sterile and clean environment with routine cleaning sessions, given the nature of the samples this room should be isolated as “a dirty room” and all molecular reagents should be kept elsewhere. Those working in this room should not enter the room in which molecular reagents are kept and laboratory clothing should remain restricted to this room.

The class II MSC should be running as ‘safe’ prior to work to ensure a stabile airflow. The cabinet should be cleaned sequentially with bleach, 80% ethanol, and RNaseZap (Sigma). The cabinet should be set up with the required reagents and waste vessels and sample bags placed directly inside class II MSC and sprayed with 80% ethanol. Barcodes are scanned for tracking and arranged into batches of ≤24 for extraction, dependent on centrifuge rotor capacity. The sample tubes, still containing the swab, should be placed into a rack, 500μl 100% ethanol (final ethanol concentration 50%) added to the tube and these are incubated at ambient temperature for 10 minutes. Top-up lysis buffer containing the internal extraction and amplification control (25 μl of 10^−3^ MS2 (~ 6 × 10^4^ PFU/ml) per 10ml of lysis buffer in this case) is next added to each sample (400μl to make 35% final ethanol concentration). The media is transferred into a spin column (NBS biologicals) over a 2ml RNA free collection tube (Thermofisher). To avoid contamination only one column is open at any one time and filter pipette tips used for each sample. The tubes should be loaded into a microcentrifuge rotor inside the class II MSC and the aerosol-tight lid closed before returning the rotor to the microcentrifuge. The samples are centrifuged for 30 seconds at 15,000 rpm (2 spins are required per sample to load the entire volume of lysis buffer).

All pass-through liquid should be discarded into designated liquid collection containers (do not mix with disinfectants containing bleach). 500μl of wash buffer 1 (1M guanidine thiocyanate in 25 mM Tris, with 10% ethanol) is added onto the columns and centrifuged for 30 seconds at 15,000 rpm. The pass-through liquid is discarded and 500μl of wash buffer 2 (25 mM Tris buffer with 70% ethanol) is added and again centrifuged for 30 seconds at 15,000 rpm. Again 500μl of wash buffer 2 is added and the tube is centrifuged for 2 minutes at 15,000 rpm, with the wash solution discarded. The silica spin column should be transferred to a new collection tube and centrifuged at 15,000 rpm for 1 minute to remove residual ethanol. The silica spin column is transferred to a new RNase free tube with the appropriate label. 100μl of nuclease free water is added to each column and left to stand for 1 minute before centrifugation for 1 minute at 15,000 rpm. The spin columns are discarded and 12μl of eluate is transferred into a 96 well plate according to ‘qRT-PCR plate layout’. The remaining nucleic acid extracts are frozen at −80°C with the location recorded on the ‘sample record’ form.

### Amplification of SARS-Cov2 nucleic acid

Once the nucleic acid (viral RNA) has been extracted, it can be amplified to detect SARS-Cov2 (see protocol 3). Notably, this work should be done in a “clean room”, and the operators should wear laboratory clothing that is restricted to this room. Movement to other working areas where biological or molecular contamination may be an issue should be restricted, and there should be no access to the dirty room.

The master mix is made up of: 12.5μl 2X Luna Universal Probe One-Step reaction mix, 0.5μl of 20pmoles/μl forward primer (ATGGGTTGGGATTATCC**T**AAATGTGA), 0.5 of μl 20pmoles/μl reverse primer (GCAGTTGT**G**GCATC**T**CC**T**GATGA**G**), 0.3μl of 10pmoles/μl MGB Probe 3 FAM (ATGCTTAG**A**AT**T**ATGGCCTC**A**C), 0.5μl of 10pmoles/μl of internal control forward primer (MS2) (supplied by Eurogentec), 0.5μl of 10pmoles/μl internal control reverse primer (MS2), 0.3μl of 10pmoles/μl internal probe (MS2 ROX), 1μl of Luna WarmStart RT Enzyme Mix (New England Biolabs) and 3.9 μl water. Once the mastermix is prepared, it can be stored 4°C short term or −20°C longer term. If using immediately, 20μl can be inoculated into a 96-well plate in clean Class II cabinet and then combined with 5μl of each RNA extract to a single well, using a different pipette tip for each well. Ideally, the master mix preparation and addition of RNA to each well should be done in separate Class II cabinets to minimize contamination.

For a negative control an extraction control containing 5μl spiked Internal extraction and amplification control (minimum of 2 wells) ae included. As an additional negative qRT-PCR control, 5μl nuclease-free water (minimum 2 wells) is included. As a positive qRT-PCR control 5μl of spiked SARS-Cov2 template plasmid is included. After adding 5μl of each sample to designated well, the plate is sealed carefully with an optically clear plastic seal using a plastic plate sealer. The plate is centrifuged for 1 minute at 1,000 rpm at 4°C and then inserted in the qRT-PCR machine (QuantStudio; Thermofisher scientific) and the run is parametrised. FAM and ROX are acquired; ROX is highlighted as the internal control, the positive and negative control wells as highlighted. The assay is run for 2 minutes at 25°C, 15 minutes at 50°C (for the reverse-transcriptase), 2 minutes at 90°C, before 45 cycles of 95°C for 3 seconds followed by 60°C for 30 seconds.

The results are determined by confirmation of the correct positive controls (amplification of the spiked target), the extraction and amplification controls of all samples (ROX channel), no amplification in the negative controls and all samples in the FAM channel with an appropriate (non-undulating or linear) sigmoidal curve equating with a CT value ≤36. The CT values of MS2 and MGB probe 3 to a Levey-Jennings plot to track quality and reproducibility of the assay [9,10].

## Results

### Establishing a workflow for SARS-Cov2 qRT-PCR

Upon the decision to rapidly establish the qRT-PCR assay we identified several challenges, these included: a) establishment and validation of a method suitable for diagnostic reporting, b) safe extraction of nucleic acid from a highly transmissible virus, c) accessing reagents required for performing extractions and amplifications, d) establishing a “clean” diagnostic workflow to minimise the risk of contamination, and e) creating a system in which HCWs could be swabbed and the data reported confidentially within a specified timeframe.

### Setting up a diagnostic qRT-PCR

In our setting, diagnosis of infections for the hospital is normally performed in the hospital diagnostic laboratory, which is co-manned by hospital and Public Health England (PHE) staff. Upon agreement with senior diagnostic staff we sought their approval that we could duplicate their in-house generated, validated assay on our equipment. The diagnostic laboratory provided access to their in-house method (designed by Martin Curran and Surendra Parmar) and provided a collection of anonymised SARS- Cov2 positive extractions (as determined by the same PCR method) and a cloned positive control. The required reagents were ordered and the QuantStudio machine calibrated to run the qRT-PCR. qRT-PCR was initially performed using existing positive samples and ten-fold dilutions of the cloned target gene (a conserved region with the ORF1 polyprotein). Upon amplification, we were able to replicate the positive signals from known positive samples (with comparable CT values of between 20 and 33) and generate a reproducible standard curve that could be used for all following amplifications and validations (Figure 1a). Additionally, during this process we validated the amplification process by the addition of a positive control; MS2 nucleic acid was added to all samples with the exception of the positive SARS-Cov2 control and the negative controls (Figure 1b). Notably, these assays were run a minimum of three occasions over differing days to assess the degree of experimental variation. Through this procedure of testing, troubleshooting, and assay development, we were able to show reproducible amplifications and have an assay ready for downstream validation.

**Figure 1.**
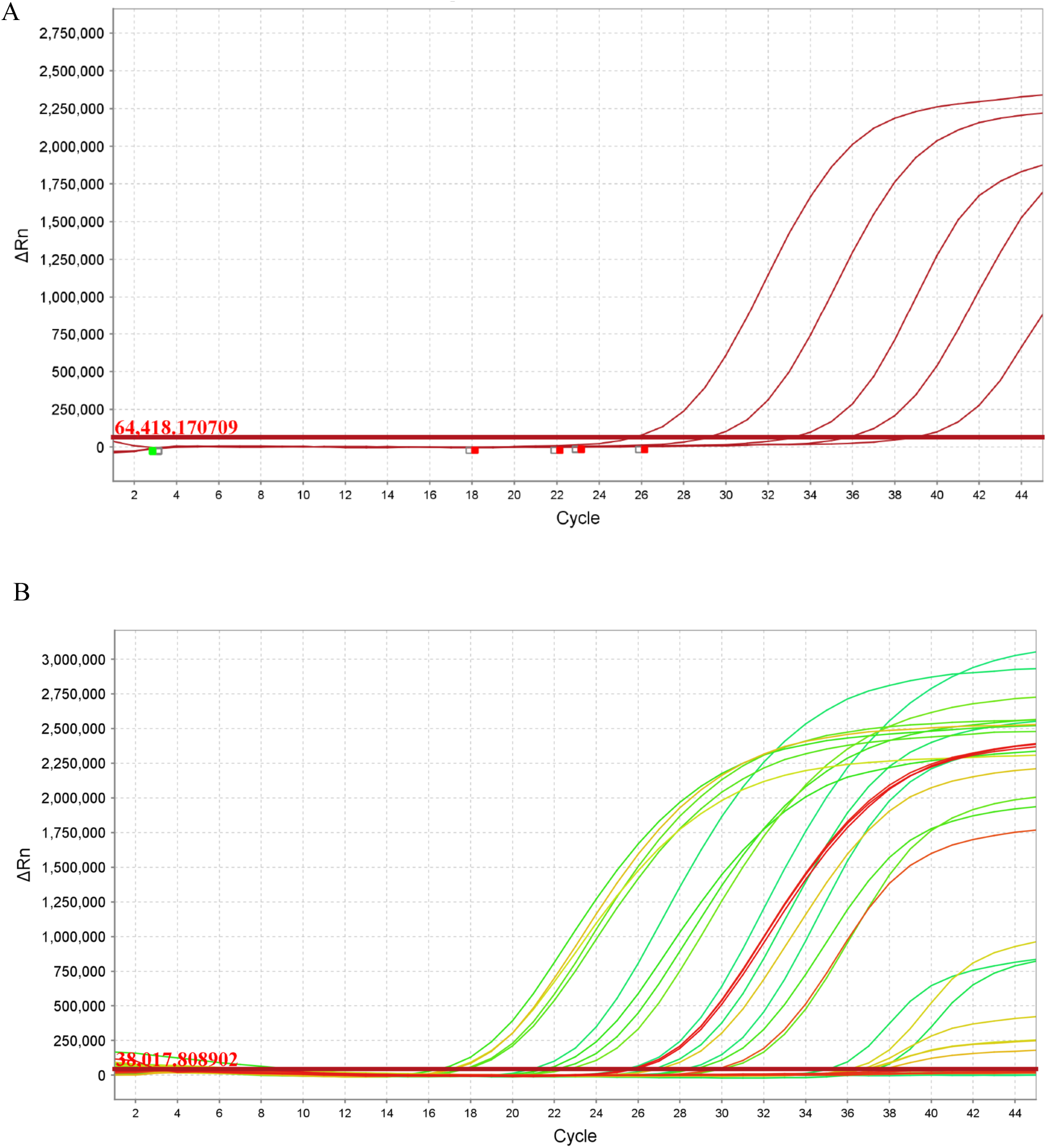
Establishing positive control qRT-PCR for SARS-Cov2 A)Amplification plot of cloned SARS-Cov2 template plasmid in 5 10-fold dilutions with FAM reporter. The x-axis displays the number of PCR cycles and the y-axis show the magnitude of normalized fluorescence signal generated by the reporter at each cycle during the PCR amplification in the form of ΔRn. B) Amplification plot of cloned MS2 control from spiked test samples with ROX reporter. The x-axis displays the number of PCR cycles and the y-axis show the magnitude of normalized fluorescence signal generated by the reporter at each cycle during the PCR amplification in the form of ΔRn. Data analysed using QuantStudio 6 and 7 Flex Realtime PCR System Software colours correspond to plate location.

### Swabbing and nucleic acid extraction

It should be noted that at the time of starting the scheme that samples with the potential to harbour SARS-Cov2 virus were classified by the UK Health and Safety Executive (HSE) as requiring a Containment Level 3 (CL3) laboratory. This level of security was required due to the infectious nature and potential for airborne transmission. Existing sampling procedures exploited a viral transport media containing ingredients to preserve the virus and restrict the growth of non-viral pathogens. The extraction of nucleic acid (or viral inactivation prior to downstream processes) in a CL3 facility was deemed a major bottleneck that could be circumvented. Consequently, we considered it essential to inactivate the sample safely at source so as to minimise risk.

A protocol was established that was risk assessed by the University Health and Safety Committee to inactivate nasal/oral swabs immediately after they are taken from the individual being sampled. The protocol is outlined in methods (protocol 2) and was established from existing methods known to chemically inactivate viruses. We utilised existing data regarding coronavirus and other highly infectious viral pathogens. Several methods, including heat inactivation were considered but the selected method using guanidine thiocyanate and β-mercaptoethanol was considered to be the most suitable for validation. Whilst existing data demonstrated that the designated approach was safe for viral extraction it had not been tested on COVID-19 patients. A recent publication has highlighted that traditional AVL lysis buffer, on which our home-made equivalent is based, does not lead to 100% inactivation of live virus when mixed at a ratio of 4:1 and left with a contact time of 10 minutes.

However, our protocol relies on the use of dry swabs, therefore any dilution effect of the lysis buffer is negated [9]. In addition, because of the locality of the testing laboratory, the minimum contact time between the swab and the lysis buffer is typically >1hr. This was followed by the additional of ethanol to a final concentration of 50% in a MSC.

### Sample workflow

A critical step in establishing a diagnostics facility is the segregation of workspace and staff, preventing the cross contamination of samples, equipment, and reagents. The mode of operation is not typical for many research laboratories where communal facilities are used according to the requirements of the specific project. The research laboratory was reorganised to create “dirty”, “clean”, and amplification areas (Figure 2a). These were strategically located in separate rooms and a strict regime was created where equipment, staff, PPE, and samples were restricted to these specific rooms. All laboratory staff were trained in the new containment structure and in the assays being performed. This component involved the transfer of materials, knowledge, and protocols between PHE Cambridge and the research laboratory. Laboratory staff were given specific roles and were restricted to the “clean” or “dirty” work areas for a single working day. Ultimately, we had developed a workflow that could be tested for screening sample from COVID-19 patients (Figure 2b)

**Figure 2.**
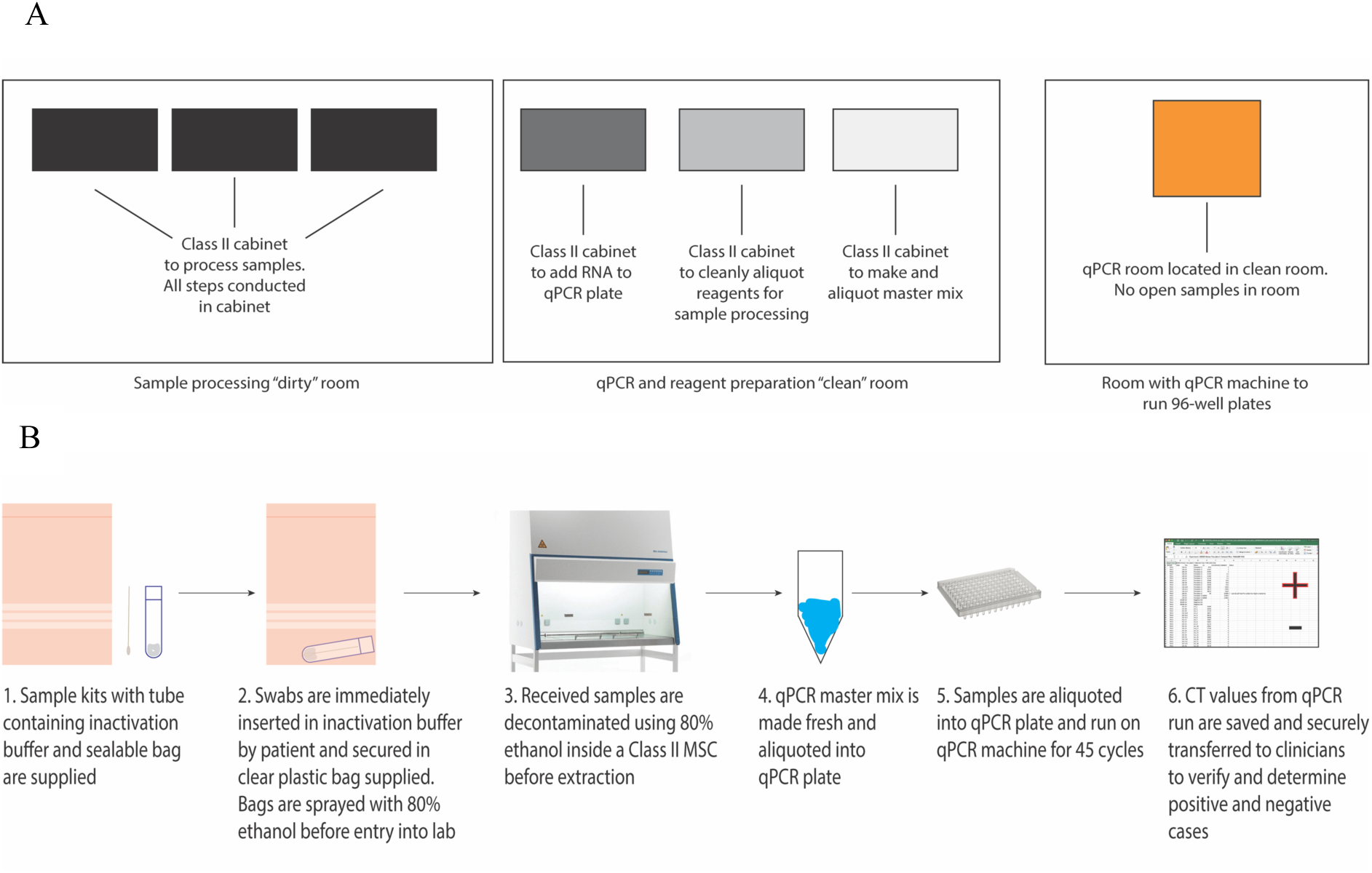
Establishing a diagnostic workflow for qRT-PCR for SARS-Cov2 A)Diagram displaying the segregation of the “dirty”, “clean” and “amplification” rooms. Note the use of separate cabinets for the preparation of reagents in the “clean” room. Individuals working in the “dirty” or “amplification” rooms are unable to enter the “clean” room on the same working day. B) Diagram showing a suitable workflow of samples from swabbing to amplification to reporting.

### Final validation of qRT-PCR assay from known COVID-19 patients

The next stage in validating the process was to run the full extraction protocol and assays on samples from patients that had previously tested positive for SARS-Cov2 in the assay performed by the hospital diagnostic laboratory. Buffers and extraction kits were constructed in the “clean” rooms and provided for patient sampling. In agreement with the hospital, for the purposes of developing a diagnostic test, we approved that a group of 20 known COVID-19 patients and a group of 20 individuals assumed not to be infected with SARS-Cov2 would be screened. Consequently, 40 swabs were taken according to the protocol from these individuals; these were dispatched to the laboratory for processing and analysis. The samples were anonymised, and research workers were blinded from knowing which samples were positive or negative. Additionally, instead of a precise 20/20 split in the provided samples, 19 were from known SARS-cov2 patients and 21 from uninfected patients, again this was not revealed to the staff performing the assay until after the tests results were known. Data from this experiment are shown in Figure 3a. There was a 100% correlation between the test results initially generated by the diagnostic laboratory and the research laboratory, with 19 generating CT values of between X and Y, and 21 generating no detectable signal. All controls were as required. At this point the assay was repeated several times to be further validated for reproducibility before being offered to the hospital for the screening of HCWs.

**Figure 3.**
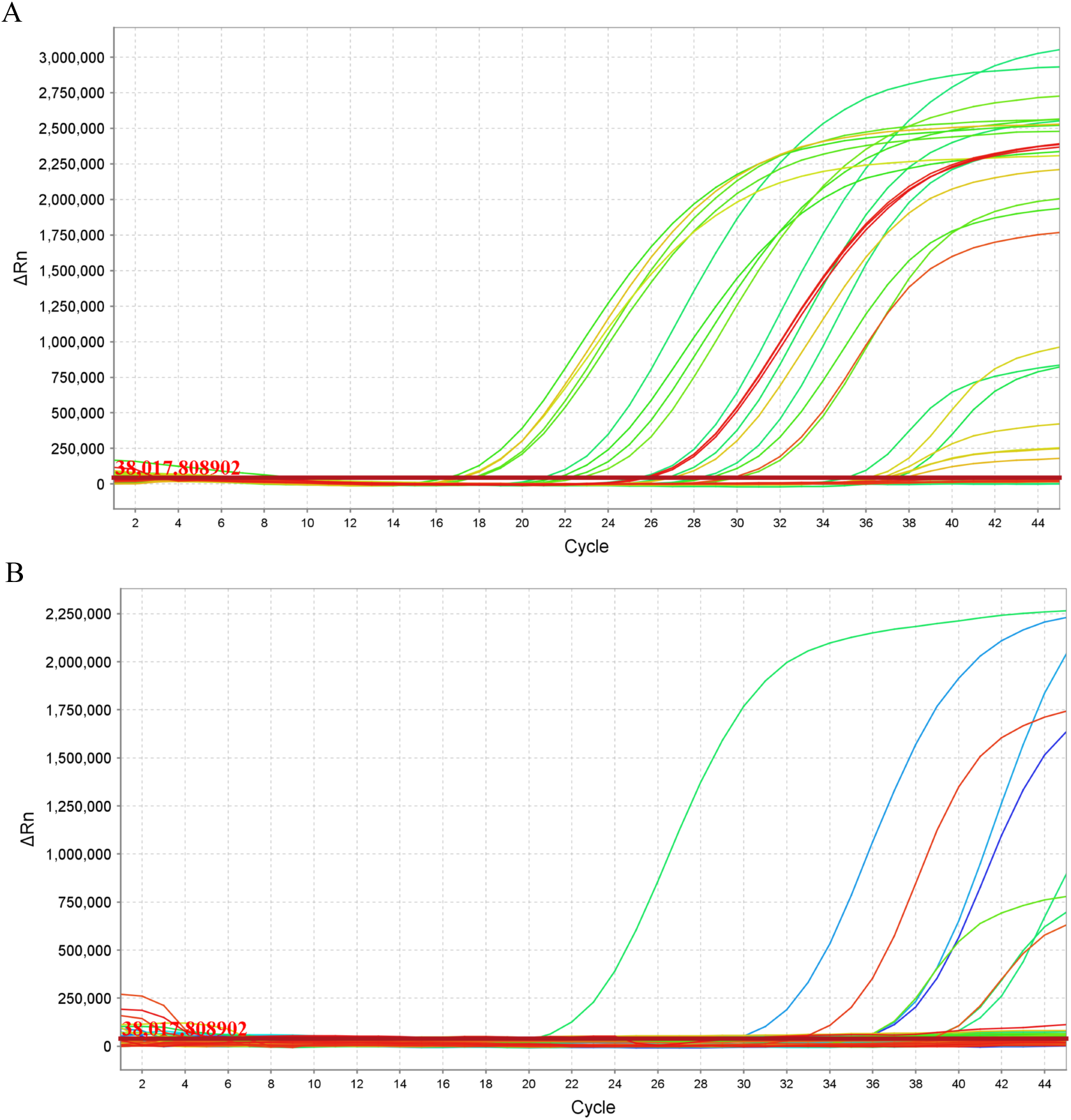
Validating and introducing a qRT-PCR for SARS-Cov2 A)Amplification plot of SARS-Cov2 from clinical samples from known COVID-19 patients. Data generated following the entire process on blinded swabs. The x-axis displays the number of PCR cycles and the y-axis show the magnitude of normalized fluorescence signal generated by the reporter at each cycle during the PCR amplification in the form of DRn. B) Amplification plot of SARS-Cov2 from samples taken from healthcare workers on first day of screening. The x-axis displays the number of PCR cycles and the y-axis show the magnitude of normalized fluorescence signal generated by the reporter at each cycle during the PCR amplification in the form of DRn. Data analysed using QuantStudio 6 and 7 Flex Realtime PCR System Software, colours correspond to plate location.

### Provision of testing for healthcare workers

Within 2 weeks of the start of the process the screening procedures were approved by the hospital and made available for hospital staff through occupational health. A firewall was built between the hospital and the research laboratory to protect confidential data without losing track of samples. A system was created where samples and data could be managed within a single system through a unique identifier number and a barcode, hence there was a logged transfer of the samples to the research laboratory, where samples could be tested, and data reported within the research laboratory. The hospital established a swabbing pod and offered structured screening to staff, and a depository for samples was established at a single point within the hospital. Samples were transported securely to the research laboratory in a risk assessed, spill proof container by selected courier, which was a member of clinical staff involved in the project. Data from the first screening run is shown in Figure 3b and permitted the detection of several positives. The CT values from these ranged from 18-36 and the turnaround time was ~4 hours from sample receival to result being available for reporting. However, given the scheduled sampling, we aimed to report the data within 24 hours.

Data were checked and validated prior to reporting by a senior member of laboratory staff. All CT values and curves were checked, and the presence of amplification in the controls was verified. All data were entered into the official hospital database and verified by a clinical virologist prior to reporting back to occupational health. Residual RNA samples were suitable for downstream analysis and we were able to contribute to ongoing COVID-19 genome sequencing projects affiliated to COG-UK.

### Troubleshooting

Clearly establishing an assay rapidly in difficult circumstances requires frequent validation, reappraisal, and troubleshooting. As the project developed more levels of management, oversight, and communication were brought in. For example, within the hospital links had to be established between those working within the wards and occupational health as the target test population were not reporting sick but were in fact being screened. Thus ethical, logistical, and practical barriers had to be identified and managed.

Within the laboratory setting, potential for the contamination of materials was a key consideration that had to be managed. For example, at one stage, background levels amplification on negative samples was elevated above acceptable levels, which was assessed to be contamination. Based on the controls used in a specific plate (having negative and positive extraction controls, swabs extracted using two different kit batches, qRT-PCR negative and positive controls), we hypothesized that the contamination was occurring in the QuantStudio equipment or sealer being used to prepare the plates. This was potentially due to SARS-cov2 DNA that was being amplified inside the machine and causing all samples to have CT values ~32. Consistency in CT value suggested that the issue was at the amplification stage and not at the extraction or qRT-PCR preparation stage.

The QuantStudio comes with a Background Calibration plate as part of its calibration plates. This plate was checked to assess whether the background profile had changed substantially. If the machine “passed” the calibration, we assessed if the profile of the background fluorescence differed from when it was previously calibrated. We performed a background calibration plate and then the bottom plate of the machine was cleaned with 10% bleach, 95% ethanol, and MilliQ water. The baseplate and optical plate were removed from the machine for deep cleaning. The baseplate was rinsed in 10% bleach, followed by MilliQ water, 95% ethanol, then water again. Liquid was aspirated from the wells, which were then wiped with a lens cleaner tissue. The optical plate above was wiped with cotton swabs containing 95% ethanol in case any dust particles were occluding the surface.

A further background calibration plate was run after cleaning and several wells were found to have high readings. To revalidate the machine, a plate of 20 mastermix plus water (negative controls) in any “problem” wells (to rule out the possibility of well-specific amplification/contamination) and additional wells scattered around the plate were assessed. Based on the location of the problem wells, contamination was often more severe at the edges of the plate, so these were also checked. The rest of the wells of the 96-well plate were filled with 10 μl of water only. The location of the “problem” wells suggested that there might have been a failed plate seal at some point, which may have released some DNA into the machine to amplify. This plate was found to give low background; several positive samples and positive and negative controls were run, and the contamination issues were found to be resolved.

## Discussion

In the continuing public health crisis, we need as much capacity as possible for supporting diagnostic services to ensure key workers and HCW are screened as frequently as possible. This places an enormous pressure on an already saturated system. The introduction of large screening services will play a huge role in tackling the epidemic in the UK and elsewhere but lacks some of the speed and flexibility that small on-site diagnostic laboratories can provide. We recognised the need to repurpose our laboratory for COVID-19 screening; this was initiated without request to provide some additional local capacity. Many academic facilities may be in a similar position but may be unsure about how to start proceedings and what regulations are in place. We suggest that groups establish the assay and processes so they can be prepared as the need arises. The route we describe here is not a scalable solution to the international lack of diagnostic testing, but a blueprint for what can be established is a standard academic research laboratory within 14 days. At the time of writing we have a full sample workflow from swabbing to diagnostic testing of HCWs at our healthcare facility, with the capacity of approximately 100 tests a day with a result provided within 24 hours. This number of tests can be expanded, but is dependent on maintaining enough extraction rooms, key staff, and of course key reagents. We sought to develop a test that works independently of kits from major suppliers, but there will potentially be issues with other resources as the crisis develops. The theoretical turnaround time is 4 hours, but this is dependent on integrating with occupational health facility and the diagnostic laboratory and ensuring there is a sustainable communication and enough staff to maintain the process.

In setting up this process there are many challenges and pitfalls, especially given the time constraints of providing a functional service that can be rapidly deployed, and we recognise that everything described here is not exhaustive. Many laboratories differ in equipment, facilities, capacity, expertise and staffing; additionally, being in close proximity to a major infectious disease centre with an excellent diagnostic facility is a major advantage. However, the methods and stages of laboratory repurposing described will, we hope, be of value to other academic laboratories in the UK and internationally that are aiming to make a useful contribution. Particularly, with some simple modifications and training we feel that this could be developed and rolled out in low and middle-income countries (LMICs), providing vital molecular capacity for this and future epidemics.

In summary, the key problems to solve are safety, reagents, cleanliness, methodology, validation, and reporting. Here, we tackled new challenges on an almost daily basis, but deactivating the virus on contact improved the process and ensured the swabs could be extracted safely in a CL2 laboratory. Access to reagents is key, and we suggest that groups become less reliant on kits from major manufacturers, unless essential. This step reduces costs and puts less pressure on existing supply chains of key kits and equipment [10]. Cleanliness is paramount, and sample flow, room segregation, and dedicated staff are essential. Having a reliable diagnostic facility that you can partner with will reduce many of the initial issues. These groups, such as the PHE laboratory here, provided excellent advice, reagents, methodology, and support for set up. Having access to good clinical facilities for validating the assay is essential; the whole process (from swab to report) needs to be comprehensively tested before being rolled out. Lastly, reporting needs to be conducted with the provision of experienced staff, again a link with clinical diagnostic facility is essential.

Here we provide a brief outline of our experience in establishing a COVID-19 diagnostic laboratory in a standard molecular bacteriology laboratory, which we hope is useful to other groups in a similar position. It was achieved under challenging circumstances through the collaborative efforts of scientists, clinical, and diagnostic staff with the ability to generate something constructive that we hope will contribute to the ongoing crisis.

## Acknowledgments

We wish to acknowledge all involved from the onset in giving their time, effort, and knowledge in getting this going. We additionally acknowledge the healthcare workers at Addenbrookes hospital, Cambridge.

---

**A blueprint for the implementation of a validated approach for the detection of SARS-Cov2 in clinical samples in academic facilities**

**Protocol 1. CoVID-19 Swabbing procedure**

**Kit contents:**

- PPE as required according to local guidelines
- Dry sterile swab in packaging
- 4ml (high sided) labelled externally threaded cryovial containing 500ul lysis buffer (double-check the name and DOB are correct)
- 80% EtOH spray for sterilising the outer tube
- Zip lock bags
- A spare collection bag and pair of gloves.

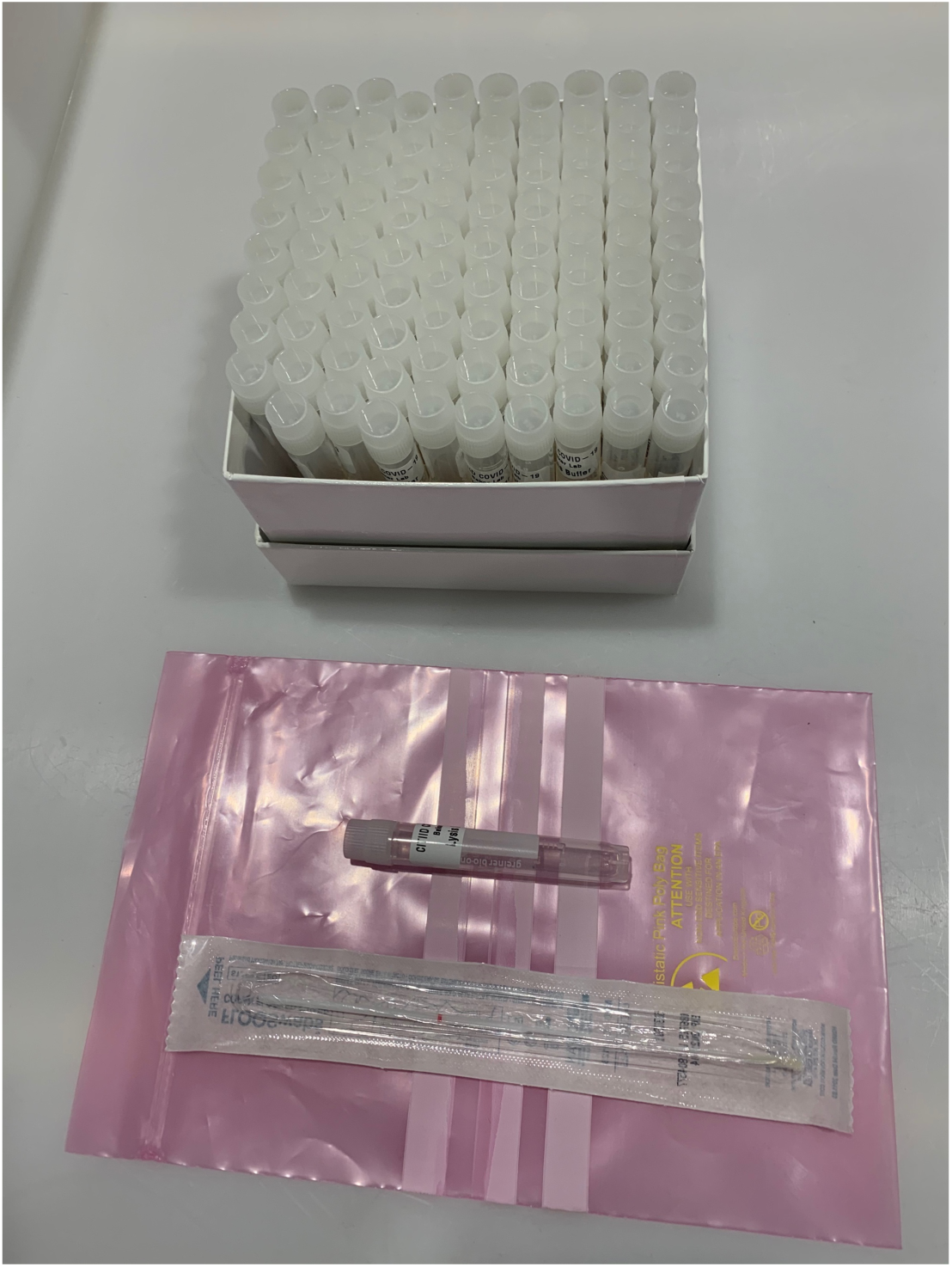

1. Check kit contents and make sure appropriate PPE is worn prior to sampling Remove swab from packet and swab (oral or nasal) the individual using the standard procedure for viral swabbing
2. Stand in front of the mirror
3. Put on the gloves
4. Remove the swab
5. **Swab** (NB one swab, two sites)
6. Open the container of lysis buffer.
7. Insert the swab. Break the swab in the container so that the swab remains in the container and the end is broken off. Seal the container tightly.
8. At arm’s length, hold out the closed swab to clinical staff so that they can spray the OUTSIDE of the swab.
9. Place the swab inside the zip lock and seal well.
10. Place the swab bag inside the designated secondary bag and seal well (i.e. so that the swab is double bagged).
11. Place the secondary bag into the designated collection box.
12. Remove gloves and dispose with this piece of paper according to local guidelines.
13. Transfer to academic laboratory and store at 4°C until extraction.

### THROAT

- Ensure you reach the very back of your throat and tonsils (i.e. not your tongue)
- Rotate the swab around this area 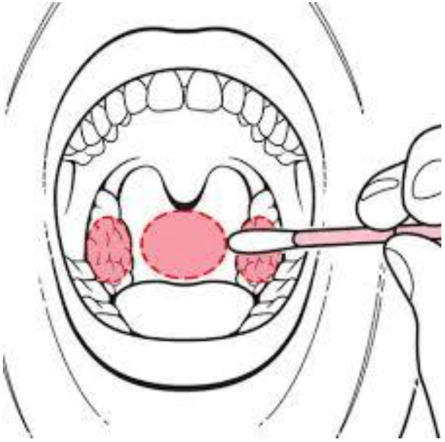

### NOSE

- Tilt your head back 70 degrees
- Enter swab along the floor of your nose (i.e. straight back, not up) until you reach resistance (this is the back of your nasopharynx)
- Rotate a few times, then withdraw slowly

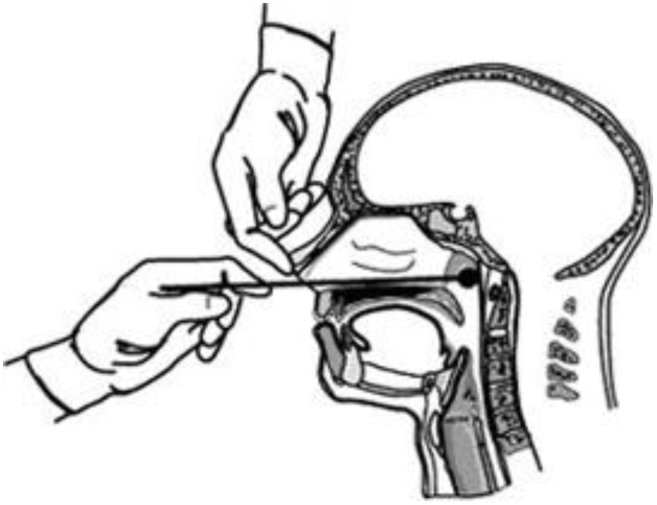

**A blueprint for the implementation of a validated approach for the detection of SARS-Cov2 in clinical samples in academic facilities**

**Protocol 2. Extraction procedure**

1. Ensure class II MSC is running as ‘safe’ prior to work to ensure a stable airflow. Clean cabinet with RNaseZap and 80% ethanol.
2. Set up class II MSC with:
  a. Pipette tips: 1ml and 200μl.
  b. waste jars: one for waste containing guanidine thiocyanate and β-mercaptoethanol and one for all other waste.
  c. container for liquid flow through collection
  d. racks for spin columns and cryovials
3. Place sample bags inside class II MSC and spray with 80% ethanol.
4. Scan barcodes into daily tracking spreadsheet and arrange into batches of ≤ 24 for extraction, depending on centrifuge capacity.
5. Open sample bag inside the class II MSC and ensure correct labelling. Spray the outside of each tube with 80% ethanol.
6. Place sample tube in a rack and add 500μl 100% ethanol (final ethanol concentration is 50%)
7. Incubate at ambient temperature for 10 minutes. To ensure complete contact inside the tube gently invert the vials several times. Label spin columns and elution tubes.
8. Add 25μl of MS2 control (at 10^−3^ concentration (~6 x 10 ^4^ PFU/ml)) per 10ml of lysis buffer (containing 0.5% β-mercaptoethanol and carrier RNA at 100 pg/μl). Label this tube ‘top-up lysis buffer’.
9. Add 400μl of top-up lysis buffer per sample (final ethanol concentration is 35%).
10. Add 600μl of sample to silica spin column. Only open one column at a time and change pipette tip between each sample. Load tubes into microcentrifuge rotor inside the class II MSC and close aerosol-tight lid before returning the rotor to the centrifuge. Centrifuge for 30 seconds at 15,000 rpm (2 spins required per sample).
11. Discard tube containing swab into designated waste jar.
12. Discard pass through liquid into designated liquid collection container. Do not mix with disinfectants containing bleach. Dispose all liquids as chemical waste.
13. Add 500μl of Wash buffer 1 to columns and centrifuge for 30 seconds at 15,000 rpm. Discard wash solution.
14. Add 500μl of Wash buffer 2 and centrifuge for 30 seconds at 15,000 rpm. Discard wash solution.
15. Add 500μl of Wash buffer 2 and centrifuge for 2 minutes at 15,000 rpm. Discard wash solution.
16. Transfer spin column to a new collection tube and centrifuge at 15,000 rpm for 1 minute to ensure all ethanol is removed.
17. Transfer spin column to a new, labelled RNase free tube.
18. Add 100μl of nuclease free water to each column and leave to stand for 1 minute.
19. Centrifuge for 1 minute at 15,000 rpm.
20. Discard spin columns and close the tubes.
21. Transfer 12μl of eluate into a 96 well plate according to qRT-PCR plate layout.
22. Freeze remaining sample at −80°C and record location on daily tracking spreadsheet.

## Reagents

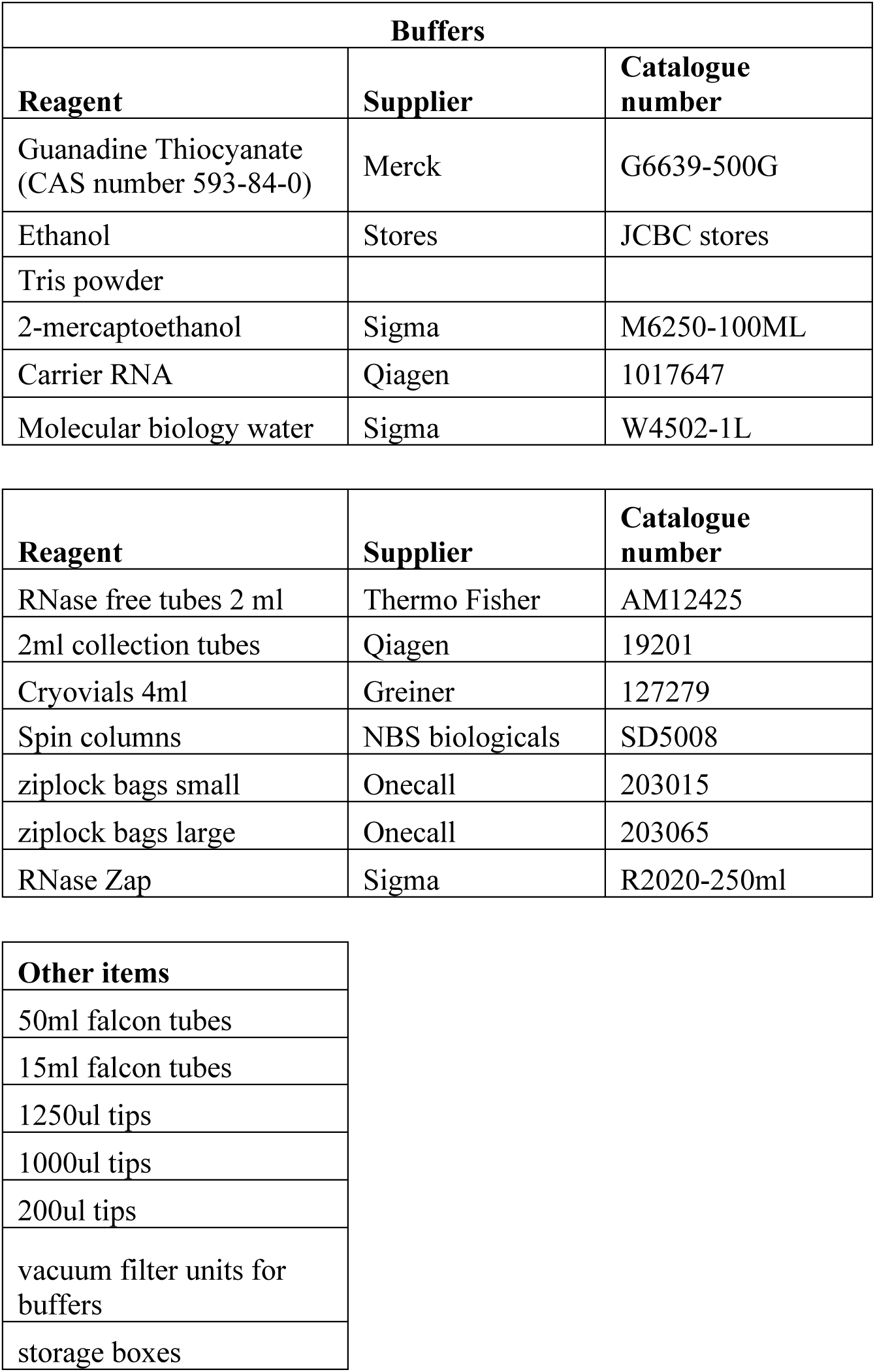

---

**Protocol 3. qRT-PCR plate setup protocol**

**Master mix** (Add in the order below):

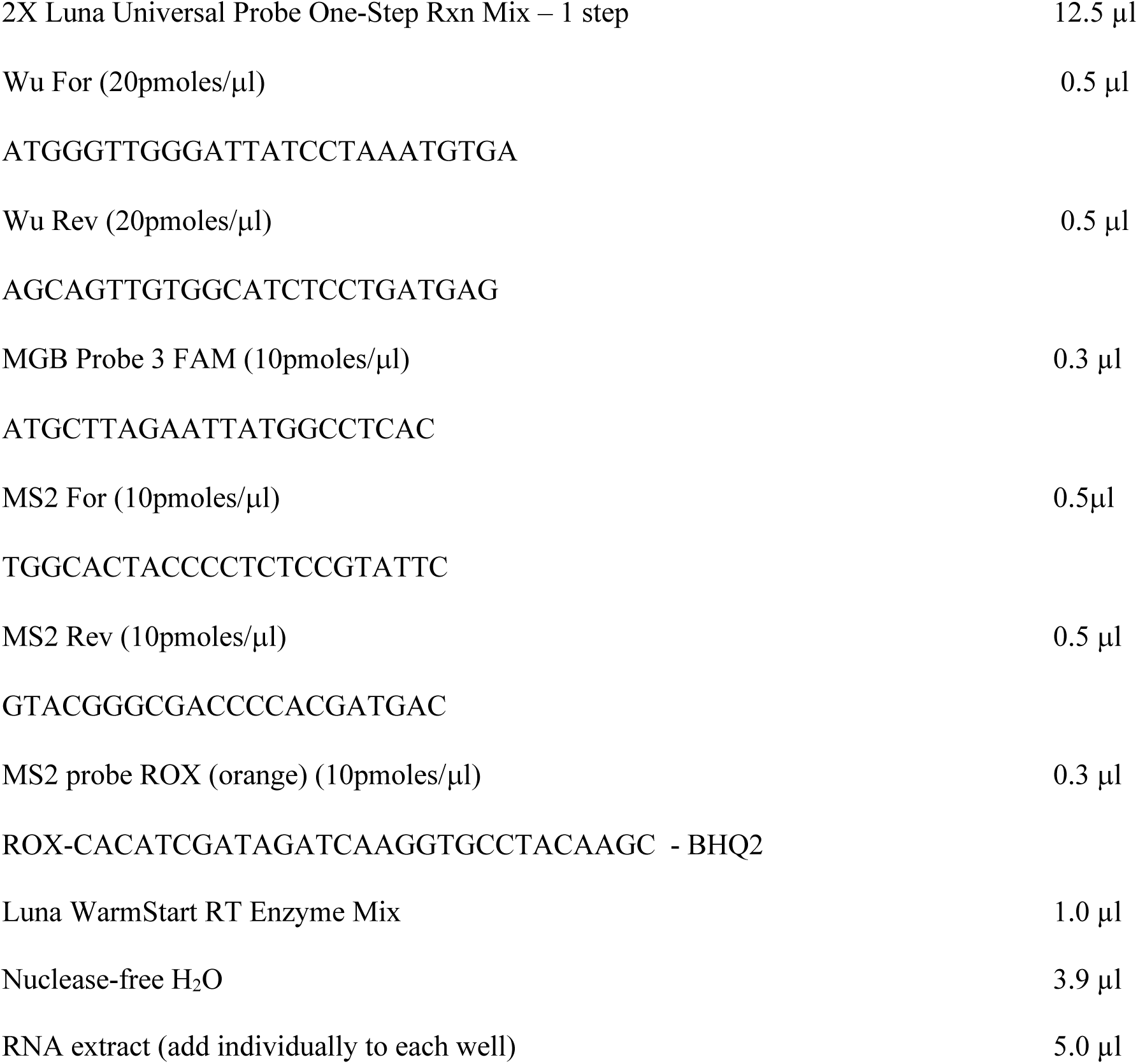

Ensure FAM and ROX are selected and calibrated.

**PCR Conditions**

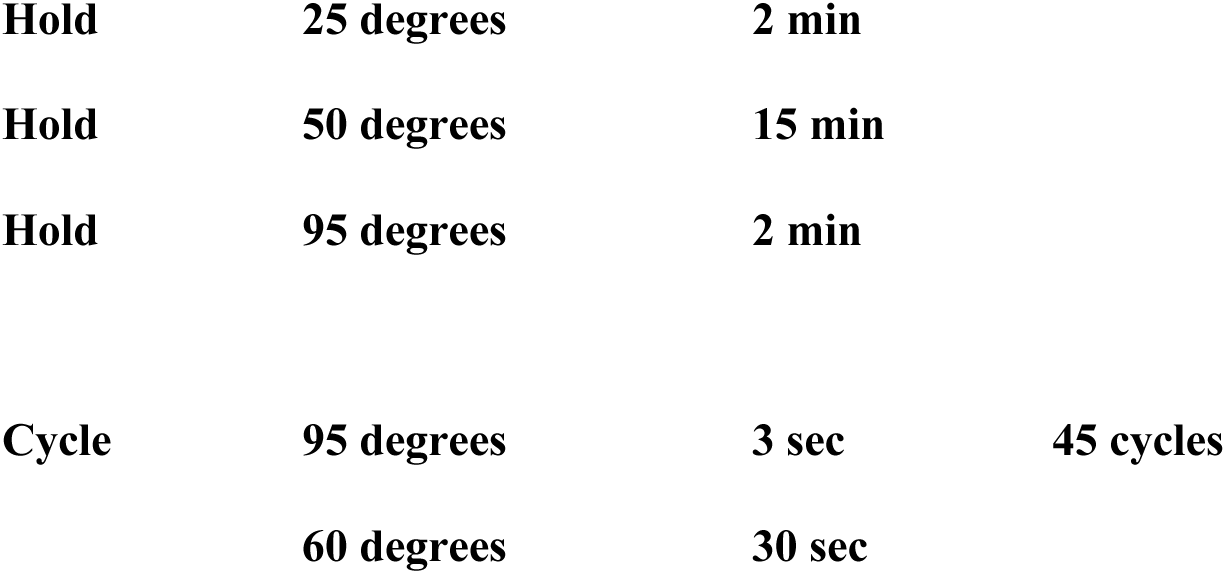

Acquire on FAM (green) and ROX (orange)

**qRT-PCR Procedure**

Once master mix is prepared, either store at 4°C short term or −20°C longer term. If using immediately, aliquot 20 μl into a 96-well plate in clean Class II cabinet.

Shift plate of mastermix to RNA-exposed clean Class II cabinet to aliquot RNA samples.

Add 5 μl of each sample to a single well, using a different pipette tip for each well.

Negative controls: extraction control containing only 5 μl spiked MS2 (minimum of 2 wells)

Negative controls: qRT-PCR control containing only 5 μl nuclease-free water (minimum 2 wells)

Positive controls: qRT-PCR control containing 5 μl spiked COVID template plasmid

For our experiments, we use: 2 extraction negative controls, 3 qRT-PCR negative controls (water), and 1 positive qRT-PCR control (COVID template plasmid).

After adding 5 μl of each sample to designated well, seal plate with plastic optically clear seal using a plastic plate sealer to avoid contamination.

Spin plate for 1 min at 1000 rpm, 4°C.

Insert plate in QuantStudio plate holder and begin run after setting up parameters:

Acquire on FAM and ROX. ROX is used as internal control. Specify control positive and negative wells.

Save and run experiment.

Export data and report results as per local guidelines.

Add CT values of MS2 and MGB probe 3 to a Levey-Jennings plot to track quality and reproducibility of the assay (Levey and Jennings, 1950; Westgard *et al.*, 1977).

**Protocol 4. Buffer Preparation**

<u>1L 25 mM Tris</u>

MW: 121.14 g/mol

Add 700-800 ml nuclease-free water to 3.02 g Tris. Spin until full dissolved.

Calibrate pH meter.

pH Tris to pH 7.0 using 10M HCl (~ 2.5 ml)

Add additional water to reach 1L volume

Steri-filter and store for use in buffers below.

<u>1L Lysis (inactivation) buffer with 4M guanidine thiocyanate</u>

Reagents:

25 mM Tris-HCl (see above)

Guanidine thiocyanate (MW: 118.16 g/mol) *handle with care

Betamercaptoethanol (0.5%)*handle with care

Carrier RNA used at 1:10000 of stock concentration (1μg/μl)

Procedure:

In a large graduated cylinder, add a stir bar, 472.64 g of guanidine thiocyanate, and 25 mM Tris to 1 L.

*Note: the guanidine takes a long time to dissolve and needs routine agitation.

Once solution is clear, decant into nuclease-free storage container.

Before use, add 0.5% betamercaptoethanol (5 ml to 1 L buffer).

Add Carrier RNA (100 μl to 1 L buffer).

Store at 4°C to retain freshness of betamercaptoethanol and carrier RNA if not using directly.

<u>1L Wash Buffer 1</u>

Reagents:

25 mM Tris buffer (see above)

1M guanidine thiocyanate (118.16 g)

100% ethanol

Procedure:

In a graduated cylinder, combine 118.16 g guanidine thiocyanate, 900 ml Tris, 100 ml 100% ethanol. Stir using a stir bar until solutes are dissolved and solution is clear. Decant into clean, nuclease-free container.

<u>1L Wash Buffer 2</u>

Reagents:

25 mM Tris buffer (see above)

100% ethanol

Procedure:

In a graduated cylinder, combine 700 ml ethanol and 300 ml Tris. Mix and decant into a clean, nuclease-free container.

---

**Risk assessment**

**RISK ASSESSMENT FOR USE OF PATHOGENIC ORGANISMS**

**In the Department of Medicine**

**Project Title:** Processing previously inactivated COVID-19 patients’ nose and throat swabs

**Investigator: Stephen Baker Laboratory Location(s):**

**Laboratory**

Human pathogens are classified by the Advisory Committee for Dangerous Pathogens into four hazard groups

(HG 1-4). The hazard group determines the containment level (CL 1-4) that is to be applied to control the risks to human health (certain pathogens and types of work e.g. clinical/diagnostics of tissue samples may be derogated from the full measures described in COSHH Regulations). HG 1 agents are essentially apathogenic, but an agent not listed in HGs 2-4 must not be assumed to be HG 1; it must be risk assessed from first principles.

All work on human pathogens must be risk assessed. Deliberate work with HG 2 pathogens and above must be notified to HSE on first use of the agent. In most cases work with potentially infected material can be conducted at CL 2 and is not required to be notified.

If the pathogen is part of a project where genetically modified forms are used, the GM risk assessment will cover all work and a separate risk assessment/HSE notification will not be required.

Note that some agents may require risk assessment/notification for the purposes of the Specified Animal Pathogens Order regulated by DEFRA.

This RA/SOP needs to be read together with Dougan Hand book Heading notes are not exhaustive.

|  |  |
| --- | --- |
| **Pathogen: Working with previously inactivated patient COVID-19 nose or throat swabs** | **HG:** |
| **Overview**<br>COSHH principles of control form a hierarchy starting with removal of the hazard, then substituting with a less hazardous form; therefore, justification for not doing so is required. | RNA extraction of previously inactivated nose and throat swabs from patients with suspected COVID-19 for sequentially qPCR.<br>COVID -19 is a new disease with limited data available at the moment, therefore during this current pandemic it is important to collect |
| **Hazard Identification in respect of Human Health**<br>Outline the pathogenic properties of the agent; routes of transmission, severity, prophylaxis/treatments available etc.<br>Who might be at risk?<br>Other lab, animal, maintenance workers?<br>Wider population?<br>Environmental stability.<br>Estimate the severity or consequence of the harmful effect were it to occur. | The main risk will be potential SARS-CoV-2 transmitted by aerosol to the laboratory worker. At the moment there is not yet a treatment or prophylaxis against the virus. To remove this risk the swabs will be directly submerged in lysis buffer (containing 4M guanidine thiocyanate, 25mM Tris pH7, 0.5% B-mercaptoethanol, and 1µl of diluted carrier RNA) at the time of sample collection. Sample collection will be performed by trained medical staff following normal precautions for clinical staff wearing the specified PPE as dictated by the ward. Submersion of the swab in the lysis buffer should be sufficient for killing virus. Additionally, to avoid handling contamination issues the tube and sample bag is to be sprayed down with 70% EtOH in the ward. Contact time >1min is effective for killing SARS-CoV-1 (Persistence of coronaviruses on inanimate surfaces and their inactivation with biocidal agents; Kampf *et al.*, 2020 Journal of Hospital Infection)<br>In the lab the sample will be unpacked inside a Class II MSC where EtOH will be added to final concentration of 50% and incubated for 10 minutes at room temp. This will render the sample safe to be used outside the cabinet.<br>CL2 workers will follow PPE rules as described in the handbook, gown and under gloves will be worn at all times<br>No virus growth or culture will be carried out at this stage and, as mentioned, all samples are processed inside the Class II MSC.<br>The worker is predominantly at risk. It would not be expected that a laboratory acquired infection would be any more severe than community acquired infection, although that risk exists if the sample is heavily contaminated.<br>In the unlikely case of the worker being exposed and subsequently developing disease there is a risk of family members being exposed as well; however, this will not be a higher risk than community exposure.<br>Any personal who are immunocompromised, pregnant, or otherwise in a higher risk group, i.e. such as pre-existing health conditions or at-risk age groups should not be processing any samples. |
| **Experimental Protocols**<br>Describe the procedures to be used and relate them to the hazards posed by the agent; titres, generating aerosols, use of sharps, infecting animals (shedding) etc. | **SOP for viral deactivation from clinical swabs prior to nucleic extraction in CL2 laboratory**<br>**Version 1.2 26<sup>th</sup> March 2020 Baker/Dougan Lab Level 5 JCBC and Goodfellow lab Level 5 Addenbrookes**<br>**Precautions**<br>• Good clinical practice and good laboratory practice to be continued at all times<br>• Ensure wearing of correct PPE (according to local guidelines)<br>• Avoid aerosolising the media in the tube during swabbing and snapping of the swab<br>• Lysis buffer contains Guanidine thiocyanate and B-mercaptoethanol (in low concentrations) so avoid leaving tube open for a prolonged period<br>• Ethanol is highly inflammable and should be kept away from heat sources<br>**Taking swabs**<br>**Required:**<br>• PPE as required according to local guidelines<br>• Dry sterile swab in packaging<br>• 4ml (high sided) cryovial containing 500µl lysis buffer (4M guanidine thiocyanate, 25mM Tris pH7, 0.5% B-mercaptoethanol, and 1µl of diluted carrier RNA)<br>• 70% EtOH spray for sterilising the outer tube<br>• Zip lock bags |
|  | 1. Make sure appropriate PPE is worn prior to sampling<br>2. Label the lysis buffer tube with the information from the sampled individual with a pen that is resistant to EtOH<br>3. Remove swab from packet and swab (oral or nasal) the individual using the standard procedure for viral swabbing<br>4. Place the swab into the provided tube containing lysis buffer and ensure the end of the swab is submerged<br>5. Snap the swab carefully to avoid disturbing the buffer<br>6. Place cap back onto tube containing the buffer and swab and ensure tight, shake to ensure even distribution of lysis buffer<br>7. Spray the outside of the cryovial with 70% EtOH until entire tube is covered<br>8. Place tube into zip lock bag and seal<br>9. Remove one glove while holding the zip lock bag in gloved hand spray the bag with 70% EtOH and pass to clean hand<br>10. Place in box for dispatch to laboratory<br>**In the laboratory**<br>**Required:**<br>• PPE as required according to local guidelines<br>• Class II MSC<br>• Zip lock bags containing swabs<br>• Microfuge<br>• 70% EtOH spray<br>• 100% ethanol<br>• Filter-tipped pipette tips and set of pipettes<br>*Inside cabinet:*<br>23. Make sure appropriate PPE is worn prior to working with the samples, lab coat and gloves should be worn at all times.<br>24. Ensure class II MSC is running as 'safe' prior to work to ensure a stable airflow<br>25. Wearing gloves and without opening the bag, check sample label and label a clean spin column outside class II MSC<br>26. Open sample bag inside the class II MSC and ensure correct labelling<br>27. Place sample tube in a rack and add 500µl 100% EtOH (final EtOH concentration is 50%)<br>28. Incubate at ambient temperature for 10 minutes with gentle agitation to ensure complete contact inside the tube<br>29. Spray down sample tubes and rack with 70% EtOH spray and remove from class II MSC<br>*Outside cabinet:*<br>30. Spin liquid through silica column in 500µl aliquots for 30 seconds at 15,000 rpm<br>31. Discard tube containing swab into clinical waste<br>32. Discard pass through liquid into suitable container. Do not mix with disinfectants containing bleach. Dispose all liquids as chemical waste.<br>33. Wash using 500µl of Wash 1 for 30 seconds at 15,000 rpm and discard wash solution<br>34. Wash using 500µl of Wash 2 for 30 seconds at 15,000 rpm and discard wash solution<br>35. Wash again 500µl of Wash 2 for 2 minutes at full speed to remove all wash solution; ensure no liquid remaining in tip of the spin column<br>36. Transfer to a fresh RNase free tube<br>37. Elute using 50µl of nuclease free water |
| **Control Measures**<br>What containment level is required?<br>Microbiological safety cabinet?<br>Any derogations?<br>Any special measures?<br>Vaccine available?<br>Training and information (shared facility?); recognition of symptoms?<br>Disinfectants.<br>Emergency procedures. | All initial sample handling has to be carried out in a **Class II MSC** unless otherwise stated in the protocol. Treat all samples as potentially infectious. For cleaning surfaces, use the approved disinfectants e.g. 70% ethanol<br>No<br>No<br>70% EtOH |
| **Environmental considerations**<br>Only use if SAPO/DEFRA relevant. |  |

Additional Notes. Last review

**Record of Individuals Carrying Out Procedure**

Procedure Risk Assessed: RNA extraction of Inactivated CoVID-19 Swaps

*“I hereby acknowledge that I have read and understood the attached risk assessment and agree to abide by and implement the control measures therein when carrying out this procedure”.*

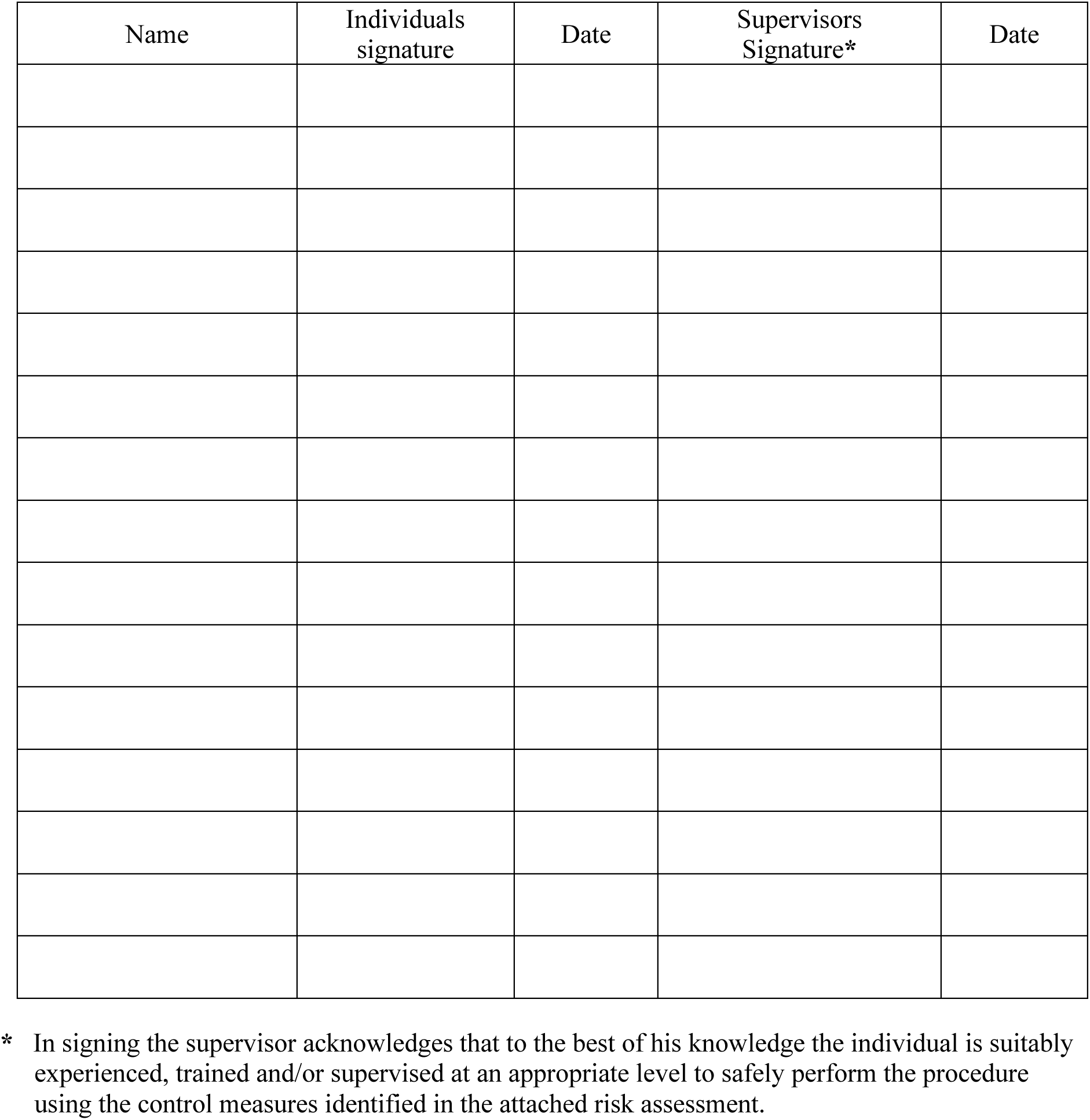

**Review Risk ALL Assessments Annually.**

Should the risk assessment be revised those persons still carrying out the procedure must sign an additional form for the revised risk assessment.

## References

1. Verity R, Okell LC, Dorigatti I, Winskill P, Whittaker C, Imai N, et al. Estimates of the severity of coronavirus disease 2019: a model-based analysis. Lancet Infect Dis. 2020;3099: 1–9. doi:10.1016/S1473-3099(20)30243-7

2. Eurosurveillance Editorial Team. Updated rapid risk assessment from ECDC on coronavirus disease 2019 (COVID-19) pandemic: increased transmission in the EU/EEA and the UK. Euro Surveill. 2020;25. doi:10.2807/1560-7917.ES.2020.25.12.2003261

3. Willan J, King AJ, Jeffery K, Bienz N. Challenges for NHS hospitals during covid-19 epidemic. The BMJ. BMJ Publishing Group; 2020. doi:10.1136/bmj.m1117

4. Bedford J, Enria D, Giesecke J, Heymann DL, Ihekweazu C, Kobinger G, et al. COVID-19: towards controlling of a pandemic. Lancet (London, England). 2020;395: 1015–1018. doi:10.1016/S0140-6736(20)30673-5

5. Tang Y-W, Schmitz JE, Persing DH, Stratton CW. The Laboratory Diagnosis of COVID-19 Infection: Current Issues and Challenges. J Clin Microbiol. 2020 [cited 14 Apr 2020]. doi:10.1128/JCM.00512-20

6. Wölfel R, Corman VM, Guggemos W, Seilmaier M, Zange S, Müller MA, et al. Virological assessment of hospitalized patients with COVID-2019. Nature. 2020; 1–10. doi:10.1038/s41586-020-2196-x

7. Bruce EA, Tighe S, Hoffman JJ, Laaguiby P, Gerrard DL, Diehl SA, et al. RT-qPCR detection of SARS-COV-2 RNA from patient nasopharyngeal swab using qiagen rneasy kits or directly via omission of an rna extraction step. bioRxiv. 2020; 2020.03.20.001008. doi:10.1101/2020.03.20.001008

8. Boom R, Sol CJA, Salimans MMM, Jansen CL, Wertheim-Van Dillen PME, Van Der Noordaa J. Rapid and simple method for purification of nucleic acids. J Clin Microbiol. 1990;28: 495–503. doi:10.1128/jcm.28.3.495-503.1990

9. Pastorino B, Touret F, Gilles M, Lamballerie X De, Remi N, Émergents V, et al. Evaluation of heating and chemical protocols for inactivating SARS-CoV-2 Méditerranée Infection), Marseille, France. Clinical samples collected in COVID-19 patients are commonly manipulated in BSL-2 laboratories for diagnostic purpose. We used the Fre. 2020; 0–8.

9. Levey S, Jennings ER. The use of control charts in the clinical laboratory. Am J Clin Pathol. 1950 Nov;20(11):1059–1066.

10. Westgard JO, Groth T, Aronsson T, Falk H, de Verdier CH. Performance characteristics of rules for internal quality control: Probabilities for false rejection and error detection. Clin Chem 1977;23:1857-

11. Newton PN, Bond KC, 53 signatories from 20 countries. COVID-19 and risks to the supply and quality of tests, drugs, and vaccines. Lancet Glob Heal. 2020 [cited 14 Apr 2020]. doi:10.1016/S2214-109X(20)30136-4

